# Engineered human induced pluripotent cells enable genetic code expansion in brain organoids

**DOI:** 10.1101/2021.06.24.449576

**Authors:** Lea S. van Husen, Anna-Maria Katsori, Birthe Meineke, Lars O. Tjernberg, Sophia Schedin-Weiss, Simon J. Elsässer

**Affiliations:** Science for Life Laboratory, Department of Medical Biochemistry and Biophysics, Division of Genome Biology, Karolinska Institutet, 17165, Stockholm, Sweden; Center for Alzheimer Research, Division of Neurogeriatrics, Department of Neurobiology, Care Sciences, and Society, Karolinska Institutet, Solna, Sweden

**Keywords:** Human induced pluripotent stem cells, Brain organoids, Genetic code expansion, Amber suppression, Non-canonical Amino acids

## Abstract

Human induced pluripotent stem cell (hiPSC) technology has revolutionized human biology. A wide range of cell types and tissue models can be derived from hiPSCs to study complex human diseases. Here, we use PiggyBac mediated transgenesis to engineer hiPSCs with an expanded genetic code. We demonstrate that genomic integration of expression cassettes for a pyrrolysyl-tRNA synthetase (PylRS), pyrrolysyl-tRNA (PylT) and the target protein of interest enables site-specific incorporation of a non-canonical amino acid (ncAA) in response to amber stop codons. Neural stem cells, neurons and brain organoids derived from the engineered hiPSCs continue to express the amber suppression machinery and produce ncAA-bearing reporter. The incorporated ncAA can serve as a minimal bioorthogonal handle for further modifications by labeling with fluorescent dyes. Site-directed ncAA mutagenesis will open a wide range of applications to probe and manipulate proteins in brain organoids and other hiPSC-derived cell types and complex tissue models.

**TOC:** 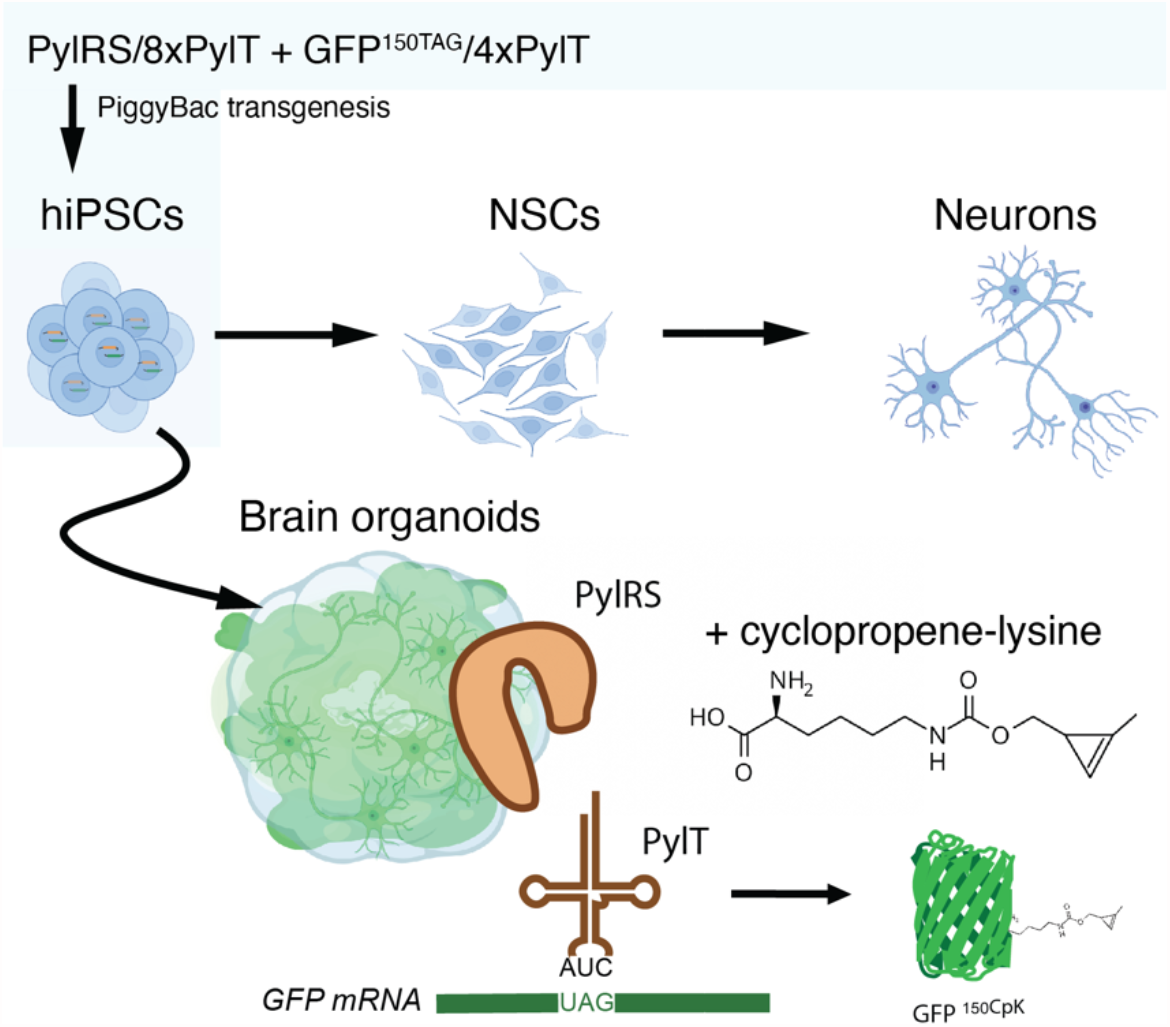

## Introduction

The systematic study of biochemical processes in neurodevelopment, neurodegeneration and other fields of human neuroscience is limited by availability of primary material and suitable model systems. Current knowledge is predominantly based on animal models or post-mortem human brain. The informative value of the prevailing rodent models is limited by the evolutionary distance between humans and rodents. ^[1]^ hiPSCs generated from patient samples and their differentiation to neurons and brain organoids could bridge the gap between animal models and clinical testing. ^[2–4]^ Since the first reprogramming of human fibroblasts to an induced pluripotent state with four defined transcription factors, OCT3/4, SOX2, KLF4, and c-MYC ^[5,6]^, robust protocols for hiPSC generation from a wide variety of patient material have been established. ^[7]^ Brain organoids recapitulate key features of the developing brain and can be derived from hiPSCs using defined protocols to model various brain regions and cell types like neurons, astrocytes and oligodendrocytes. ^[8–10]^ Cerebral organoids have been successfully used to model neurological diseases, such as Alzheimer disease ^[11]^, Parkinson disease ^[12]^, microcephaly, autism spectrum disorders and Down syndrome. ^[13]^

Non-canonical amino acids (ncAAs) can introduce chemical functionalities not found in nature into a protein of interest. Genetic code expansion towards ncAAs requires a tRNA and an aminoacyl-tRNA synthetase (aaRS) both orthogonal to the host cell (i.e. not interacting with the host translational machinery). The pyrrolysyl aaRS/tRNA pair (PylS/PylT) from methanogenic archea is routinely used to suppress amber (UAG) stop codons and introduce a ncAA in response. This approach, termed amber suppression, has been used successfully in bacteria, yeast, mammalian cell culture, and animal models. ^[14–18]^

The ncAAs introduced via genetic code expansion cover a wide repertoire of chemical groups, including bioorthogonal handles, crosslinkable moieties and photocages. ^[19]^ ncAAs have been shown to be invaluable tools for studying proteins important for neurobiological processes and pathophysiology, such as G-protein coupled receptors ^[20–23]^ and ion channels. ^[24,25]^ ncAAs are particularly useful in applications where the protein under study cannot or should not be modified in a significant manner by larger protein modifications, such as fluorescent protein fusions or affinity tags. As an example, a ncAA has enabled fluorescent labeling of the 42 amino acid long amyloid ß-peptide fragment implicated in Alzheimer disease. ^[26]^

Genetic code expansion has been implemented in rat or mouse neurons through a variety of strategies. ^[24,27]^ Using viral delivery, electroporation or lipofection, tRNA-Synthetase/tRNA pairs were also introduced transiently into mouse brains or brain slices. ^[28,29]^ However, no universal and efficient approaches exist to expand the genetic code of hiPSC-derived human cultured neurons or across entire brain organoids. Here we report the generation of hiPSCs with an expanded genetic code, enabling stable and efficient amber suppression in hiPSCs and neurons, as well as cerebral organoids derived from hiPSCs through in vitro differentiation. Genetic code expansion in cerebral organoids will facilitate implementation of the wide variety of ncAA-based technologies developed for mammalian cells to relevant human model systems for studying neurological diseases.

## Results and Discussion

Stable genetic code expansion has been previously achieved in mouse embryonic stem cells using PiggyBac-mediated transgenesis. ^[18]^ We rationalized that establishing the PylRS/PylT system in hiPSCs would provide a route to ncAA incorporation via amber suppression in a wide variety of cell types that can be differentiated from hiPSCs. Below, we describe the generation of hiPSCs line CTL07-II-AS with an expanded genetic code (Figure 1) from the hiPSC line CTL07-II. ^[30]^ CTL07-II-AS hiPSCs were differentiated into neural stem cells (NSC), neurons and cerebral organoids. We further demonstrate that amber suppression activities are maintained in mature neurons and neuronal organoid tissue.

**Figure 1.**
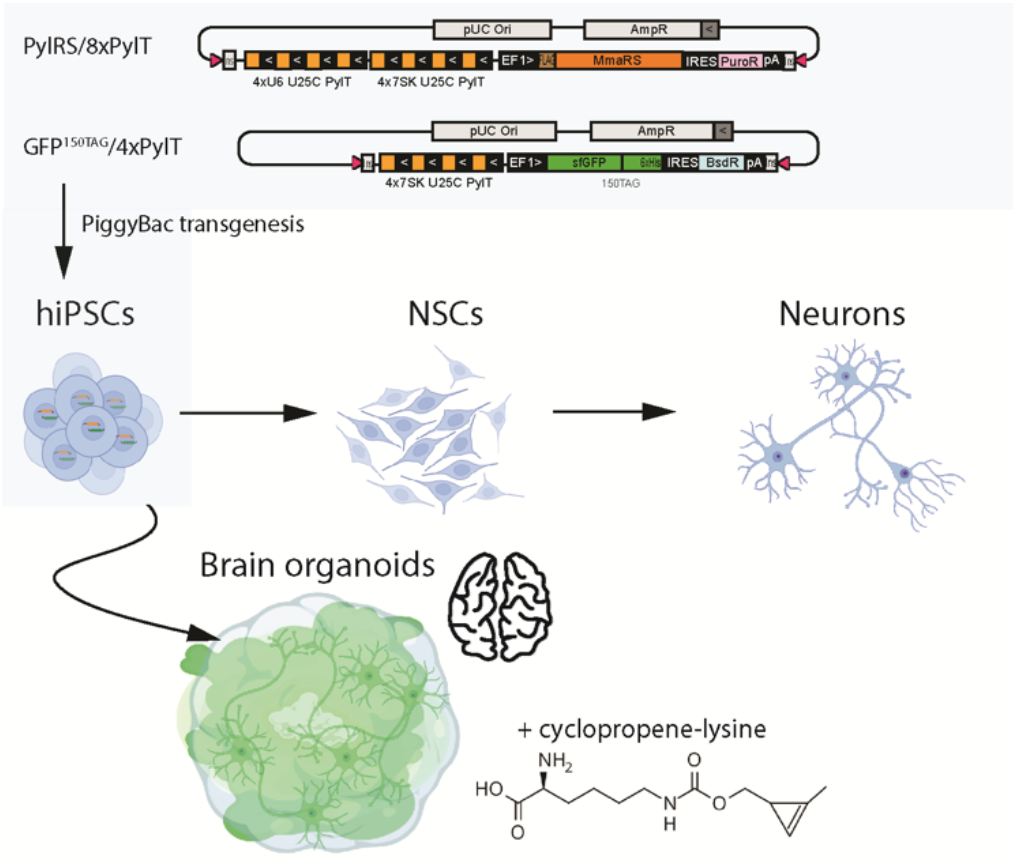
Integration of amber suppression machinery in hiPSCs enables derivation of neurons and brain organoids with an expanded genetic code. Cells are cotransfected with a PylRS expression vector and a PylT/sfGFP^150TAG^ reporter plasmid, carrying a repeat cassette for PylT expression, the PylS gene to produce PylRS and an amber stop-codon containing GFP gene. hiPSCs are differentiated to NSCs, neurons and cerebral organoids. Culturing hiPSC, NSCs, neurons or organoids in the presence of 0.2 mM cycloproprene-lysine (CpK) leads to suppression of the amber stop codon and the production of GFP with a site-specifically incorporated CpK moiety that can be subsequently reacted using biorthogonal chemistry.

### PiggyBac-mediated integration of the orthogonal PylRS/tRNA pair

To generate hiPSCs with an expanded genetic code, we used an updated two-plasmid system for integrating amber suppression machinery and reporter ^[18,31]^ using PiggyBac transposase (PBase). In the two targeting plasmids, expression cassettes include four or eight tandem repeats of PylT and are flanked by inverted repeats for transposition (Figure 1). The 8xPylT/PylS expression construct encodes FLAG-tagged Methanosarcia mazei PylS and puromycin resistance genes, and a total of eight PylT genes (four tandem repeats with U6 promoter and four tandem repeats with h7SK promoter). The 4xPylT/sfGFP^150TAG^ reporter plasmid encodes for GFP protein with an amber codon at position 150 (GFP^150TAG^), carries four tandem repeats of h7SK-PylT and a blasticidin resistance gene (Figure 1). Since transfection efficiency via lipofection in hiPSCs is low, the 8xPylT/PylS plasmid was integrated in a first co-transfection with PBase and stable transfectants were selected with puromycin. Then the 4xPylT/GFP^150TAG^ reporter plasmid was co-transfected with PBase and double integrants were selected with puromycin and blasticidin, resulting in the polyclonal cell line CTL07-II-AS (Figure 2A).

**Figure 2.**
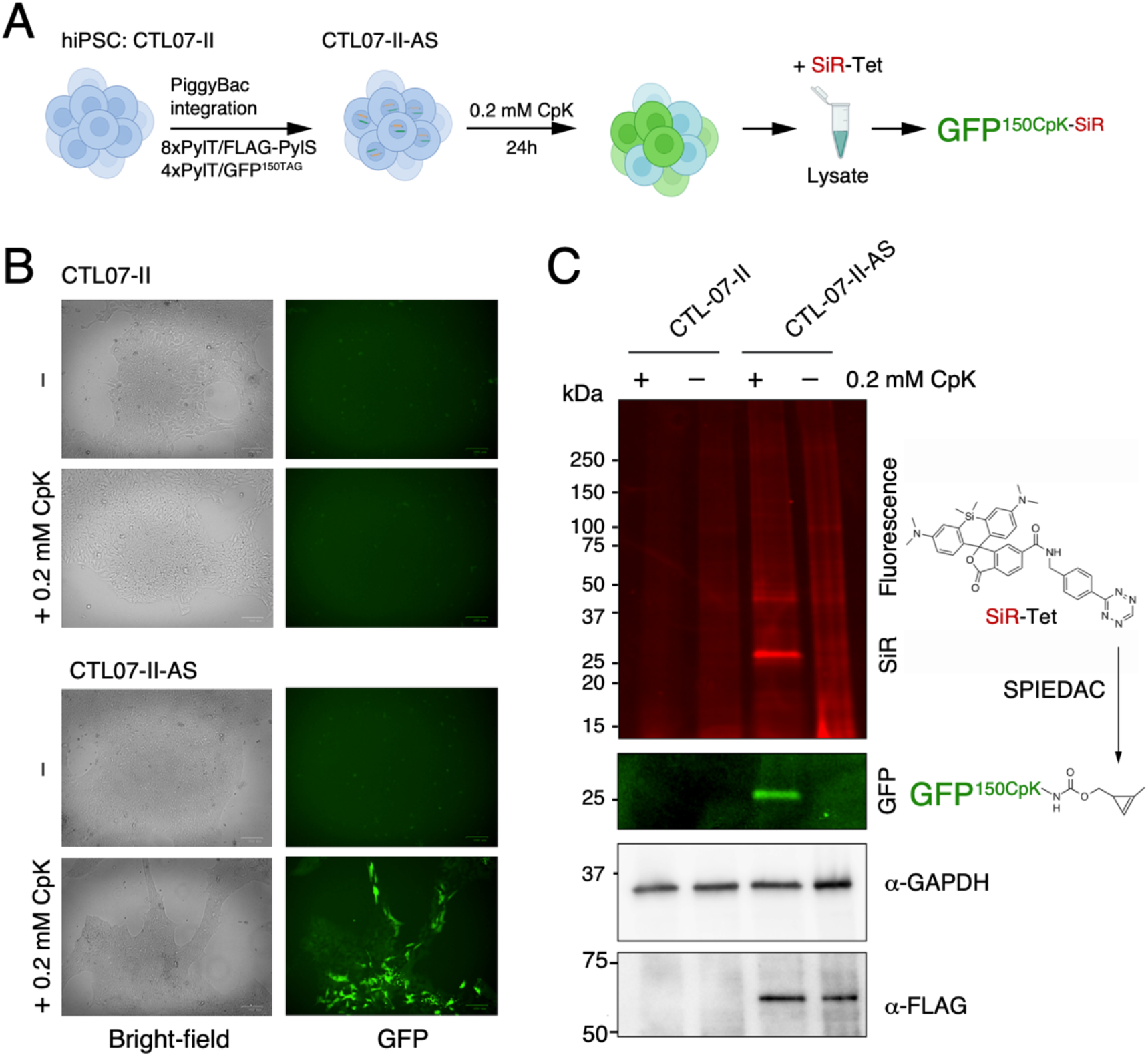
Stable amber suppression in hiPSCs. A) Scheme for generating CTL07-II-AS, a hiPSC line with expanded genetic code, CpK incorporation in the GFP^150TAG^ reporter and subsequent SPIEDAC fluorescent labeling with SiR-Tetrazine. B) Brightfield and green fluorescent images of CTL07-II and CTL07-II-AS. CTL07-II-AS cells that were incubated with 0.2 mM CpK for 24 h before imaging express GFP, while cells cultured without CpK and the parental cells do not show any fluorescence. C) SDS-PAGE and fluorescent imaging of cell lysates of CTL07-II and CTL07-II-AS cultured in the presence or absence of 0.2 mM CpK for 24 h incubated with 1 µM SiR-Tetrazine to perform SPIEDAC labeling of CpK-bearing proteins. Western blot shows FLAG=PylRS expression in CTL07-II-AS cells. GAPDH was used as loading control.

### Efficient and selective non-canonical amino acid incorporation in hiPSCs

To validate the generation of hiPSCs with an expanded genetic code, we incubated CTL07-II-AS cells with 0.2 mM cyclopropene lysine (CpK) for 24h. GFP was expressed in the CTL07-II-AS cells only in the presence of CpK (Figure 2B), indicating that the GFP^150TAG^ stop codon is efficiently suppressed by the PylRS/PylT system. We also confirmed the expression of FLAG-PylRS in the CTL07-II-AS by western blot (Figure 2C). GFP levels were heterogeneous (Figure 2B), indicating that not all the cells in the population had strong amber suppression activity. Incorporation of CpK at GFP^150TAG^ was confirmed by performing SPIEDAC labeling of the cyclopropene moiety with *silicon* rhodamine (SiR)-Tetrazine in cell lysate, subsequent SDS-PAGE and fluorescent imaging (Figure 2C). A prominent band in the SiR channel overlapped with the fluorescent GFP band (Figure 2C, S1). Notably, the low abundance of background bands in the SDS-PAGE after SiR labeling demonstrates that the amber codon is selectively suppressed in CTL07-II-AS despite the presence of amber stop codons in many endogenous genes (Figure 2C). While this result hinted that the proteome of CTL07-II-AS hiPSCs was not collaterally affected by the amber suppression machinery, we also confirmed that expression of the pluripotency marker OCT4 was not affected by 24h incubation with 0.2 mM CpK (Figure S2). Nevertheless, we note that applications of genetic code expansion in hiPSCs or derived cell lines need to consider inference from off-target effects of amber suppression activity, and it will be important to optimize concentration and incubation time of ncAA to minimize such effects.

### Differentiation of hiPSCs with an expanded genetic code

hiPSCs can be differentiated to neural stem cells (NSCs), a stable population of multipotent and self-renewing progenitor cells that can be further differentiated to neurons, astrocytes and oligodendrocytes ^[32]^ (Figure 3A). Differentiation to NSCs was performed using serum free neural induction medium within one week. Further differentiation to neurons was performed by removing bFGF and EGF stem cell growth factors and increasing B27 for three additional weeks (Figure 3A). We followed the appropriate course of differentiation to mature neurons by immunofluorescent staining for the marker proteins OCT4, SOX2, NESTIN and MAP2 (Figure 3B). OCT4 is a marker for pluripotent stem cells whereas SOX2 is expressed throughout pluripotent and neural progenitors (Figure 3C). NESTIN is expressed in neural progenitors but not in mature neurons, while MAP2 is also expressed in mature neurons (Figure 3C). Hence, we could confirm the successful differentiation of hiPSCs with an expanded genetic code into neurons.

**Figure 3.**
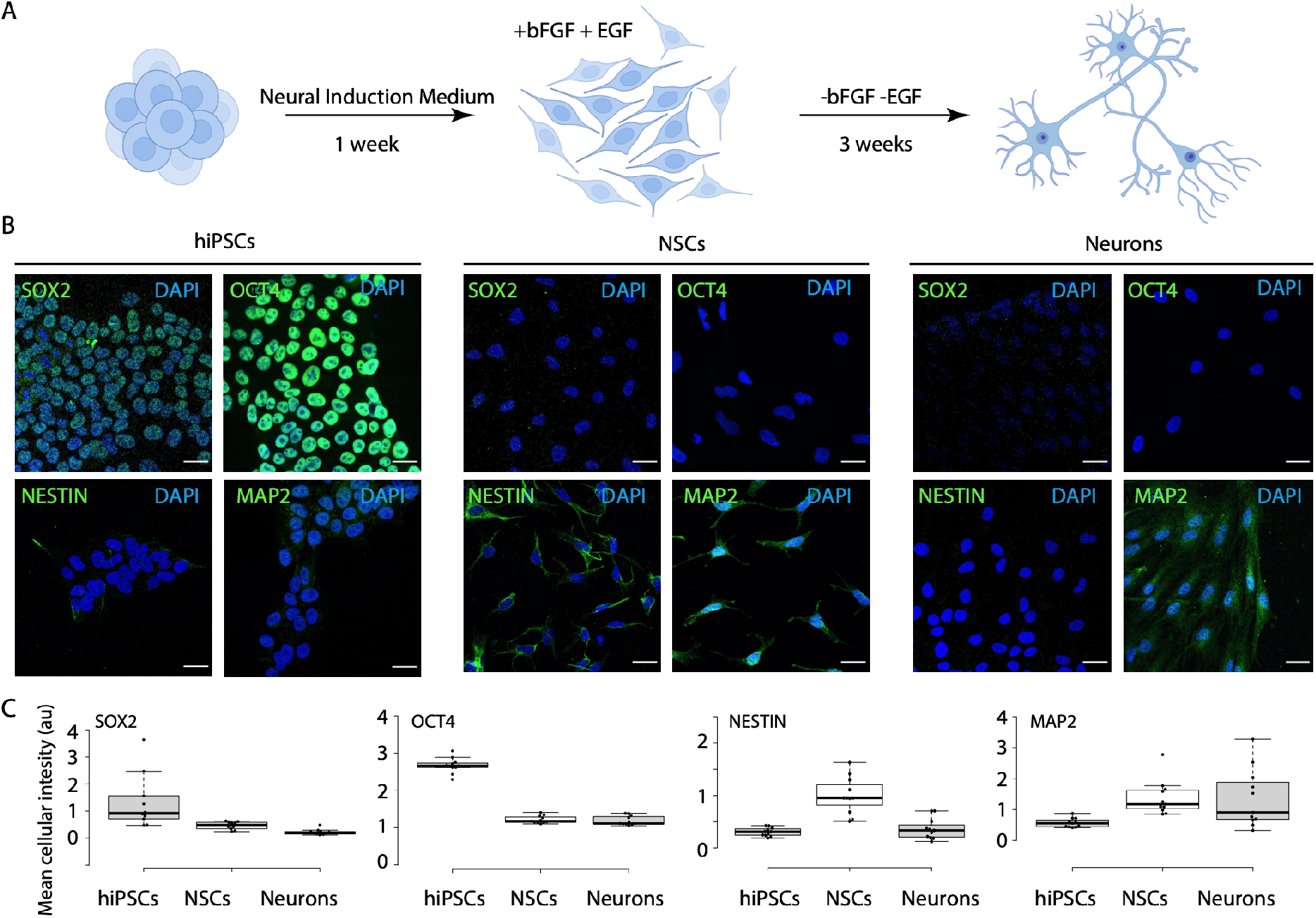
Neuronal differentiation of CTL07-II-AS. A) Scheme for differentiation of hiPSCs to NSCs and neurons. B) Immunofluorescence showing expression of stem cell markers SOX2 and OCT4 in hiPSCs, NSC marker NESTIN in NSCs and neuronal marker MAP2 in 3 week old neurons. Scale bars correspond to 30 µm C) Image quantification determining mean fluorescence intensity in each image (n=11) divided by the number of cells.

### Amber suppression in NSCs and neurons

Genomic integration of the amber suppression machinery in pluripotent cells should in principle allow amber suppression in all derived cells and tissues. The EF1 promoter used here to drive mRNA expression and h7SK promoter for PylT are thought to be active in a cell-type independent manner. While a 24h incubation with 0.2 mM CpK was sufficient to elicit robust GFP^150TAG^ expression in hiPSC, even an extended incubation with CpK for 7 days elicited an intermediate GFP fluorescence in neurons and only low GFP signal in NSCs (Figure 4A, B). The expression of GFP was heterogeneous within the population in differentiated cells, as expected from the heterogeneous hiPSC starting population (Figure 4A, B). No GFP was produced in the absence of CpK in any cell type (Figure 4B, S3). These results demonstrate that stable incorporation of the PylS/PylT pair in hiPSCs enables genetic code expansion in derived terminally differentiated cells. Incorporation of CpK in neurons was further validated by labeling live neurons with SiR-Tetrazine dye after incubation with 0.2 mM CpK for 7 days (Figure S4). Overall, amber suppression efficiency appears to vary across different cell types, which could be a function of promoter strength for PylS and PylT genes, as well as the efficiency of translation termination competing with amber suppression in the respective cell type. Further, epigenetic silencing may lower transgene expression over time. Still, neurons had higher average GFP fluorescence than NSCs (Figure 4B), potentially owing to the fact that NSCs still divide and thus the GFP production is diluted continuously, whereas it can accumulate to higher levels in post-mitotic neurons in the 7-day incubation period with ncAA.

**Figure 4.**
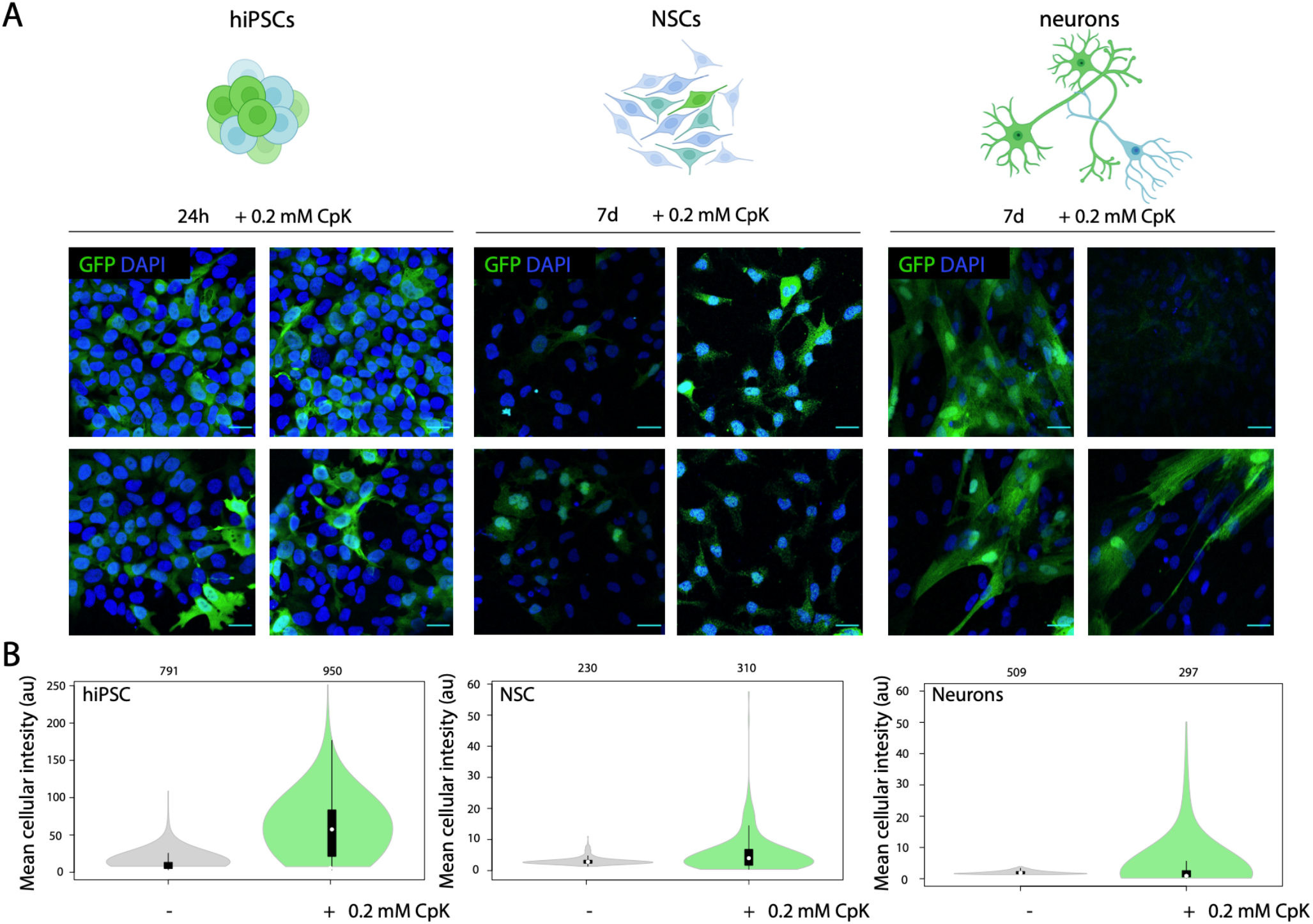
*In vitro* differentiation enables amber suppression in neurons. A) Four representative images from immunofluorescence microscopy of CTL07-II-AS hiPSCs, derived NSCs and neurons showing expression of GFP (green) and DAPI (blue). hiPSCs were cultured in the presence of 0.2 mM CpK for 24 h. NSCs and neurons were cultured in the presence of 0.2 mM CpK for 7 days. Scale bars correspond to 30 µm B) Image quantification showing the mean fluorescence intensity in each cell.

### Genetic code expansion in cerebral organoids

Having demonstrated amber suppression activity in terminally differentiated neurons from the stably integrated transgenes, we sought to derive more complex tissues from the CTL07-II-AS hiPSC line. Genetic code expansion in cerebral organoids would be even more relevant for enabling site-specific ncAA mutagenesis in studies of neurodevelopment or neurodegenerative diseases. For example, polygenic diseases like autism spectrum disorders affect multiple cell types and their connectivity and cooperativity in the brain [10].

To this end, we differentiate CTL07-II-AS hiPSCs to cerebral organoids over the course of a 40-day protocol, by first promoting the formation of embryoid bodies, which we differentiated further to neuroepithelial organoid and finally to mature organoids with cortical-like regions (Figure 5A). The successful differentiation was confirmed by immunofluorescence microscopy of fixed organoid slices, in which we identified ventricular progenitor zones characterized by SOX2 expression and MAP2 neurons in the periphery (Figure S5). CTIP2, a deep-layer subcortical projection neuron marker was expressed in the outer layer of the organoids (Figure S5). We imaged live organoids which were grown in the presence or absence of 0.2 mM CpK for 14 days on a Zeiss Lightsheet Z.1 microscope (Figure 5A). We observed strong but heterogeneous GFP fluorescence in the presence of CpK across the organoid (Figure 5B, Movie 1). The particularly high GFP fluorescence in neuron-like cells, and the lower fluorescence in neighboring cells made it possible to observe the neuronal morphology pervading the three-dimensional organoid structure (Figure 5C, Movie 2,3). We subsequently fixed the organoids for cryosectioning and immunofluorescence staining, and also observed the highest GFP intensity in neurites and soma of neurons (Figure 5D). The distinctive labeling of neurons was surprising considering that no neuron-specific promoter was used, but consistent with our observation that cultured neurons accumulated more GFP than neural stem cells. Where we were able to image more internal regions, we also observed strong GFP fluorescence in tightly clustered cells forming luminal structures akin neuroepithelial rosettes (Figure S6, Movie 4,5). In summary, these data demonstrate that genetic code expansion is possible in cerebral organoids with particularly high efficiency in differentiated neurons and luminal cells. The non-canonical acid diffuses sufficiently well into the organoid to elicit efficient amber suppression also in deeper layers. Notably, genetic manipulation of mature organoids is difficult since viruses, lipofection or electroporation delivers DNA predominantly to the outermost layers. Thus, deriving organoids from hiPSCs with a stably expanded genetic code represents a key advantage in achieving amber suppression across the entire organoid. As discussed above, amber suppression efficiencies appeared to greatly vary with cell type, suggesting that amber suppression may be modulated by cell type-specific properties, such as promoter strength and termination efficiency.

**Figure 5.**
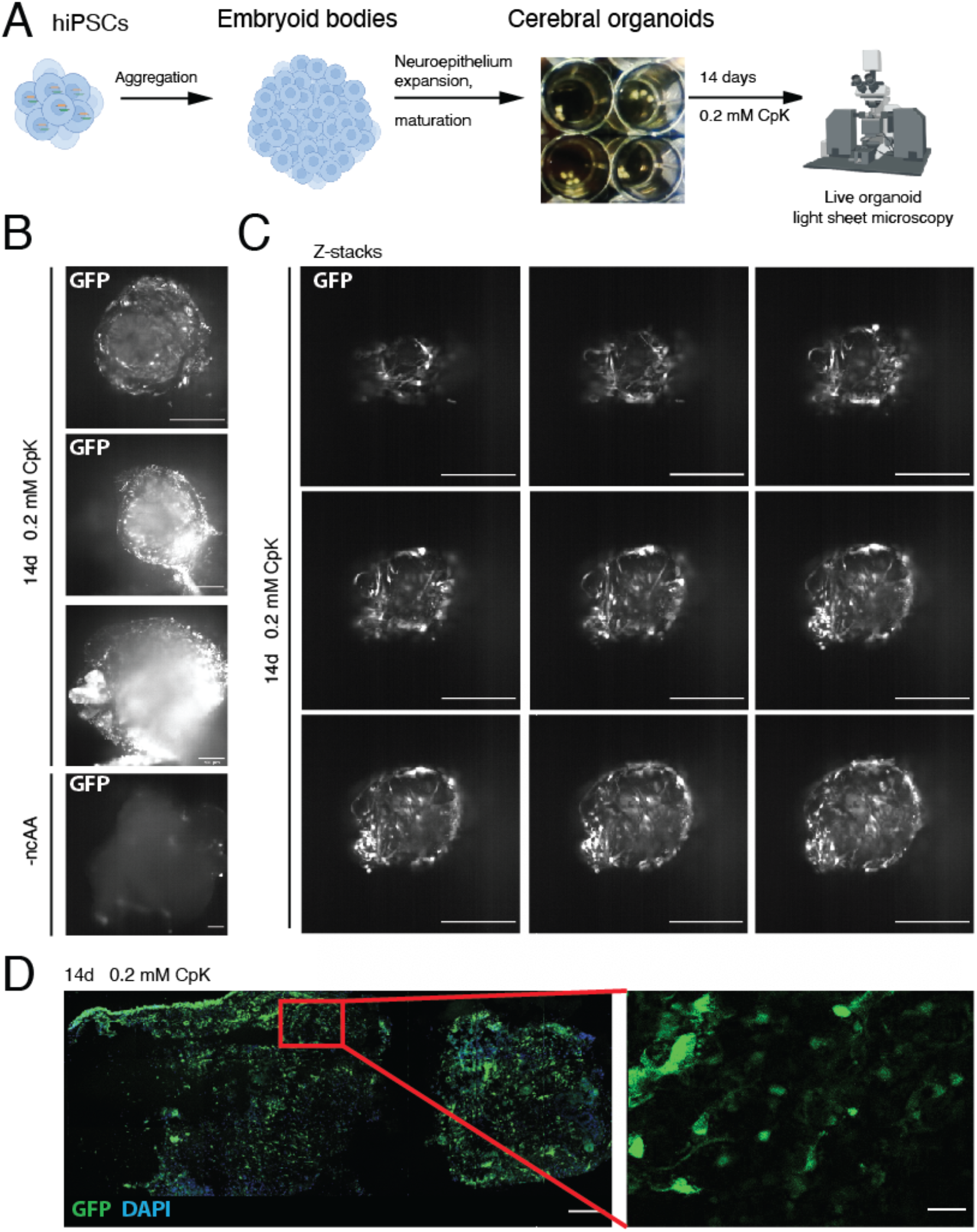
Stable amber suppression in cerebral organoids. A) Scheme for differentiation of hiPSCs into cerebral organoids. The organoids were differentiated for 40 days, cultured in the presence of 0.2 mM CpK for 14 days and imaged via light-sheet microscopy. B) Live cell light-sheet microscopy of cerebral organoids to detect GFP expression in organoids cultured in the absence or presence of 0.2 mM CpK. Selected planes are shown. Scale bars correspond to 250 µm. See Movie 1 for entire Z-stack C) Ligh-sheet microscopy Z-stack images through the tip of the cerebral organoid cultured with 0.2 mM CpK. 250 µm scale bar. See Movie 2 and 3 for corresponding Z-stack and 3D reconstruction. D) Immunofluorescent staining of fixed and cryosectioned organoid with a GFP antibody (green). DAPI is shown in blue. Scale bars correspond to 30 µm.

In summary, we have created a hiPSC line with an expanded genetic code and demonstrated that genomically integrating the PylRS/PylT pair at the hiPSC stage allows derivatization of differentiated cells and complex tissues with an expanded genetic code. We anticipate that our approach of engineering hiPSCs will enable to generate many other cell types and tissues with expanded genetic code through established differentiation protocols. We envision that the choice and optimization of promoters will be crucial to maximize ncAA incorporation efficiency in the desired cell type. For example, tissue-specific promoters for the PylS gene could be exploited for restricting PylRS expression and thus amber suppression activity to a specific cell type or lineage. Our modular transgenesis approach also leaves the opportunity to generate hiPSCs and derived cells or tissues stably expressing PylRS/PylT, combined with an alternative delivery of the protein of interest e.g. through lipofection or viral transduction.

We believe that human brain organoids with an expanded genetic code will provide a unique platform to study molecular mechanisms by using ncAAs to probe and manipulate proteins.

## Acknowledgements

We acknowledge support from the Light Sheet Microscopy Pilot Facility at SciLifeLab and the National Microscopy Infrastructure, NMI (VR-RFI 2019-00217) for providing access and support with light sheet microscopy. We acknowledge the iPS Core Facility for providing the CTL-07-II hiPS cell line. Illustrations have been made with BioRender.com. For financial support, we acknowledge Stiftelsen för Gamla Tjänarinnor (LSvH), Thomas Olausson Stiftelse (LSvH), and the Gun and Bertil Stohnes Foundation (LSvH), Ming Wai Lau Center for Reparative Medicine (SJE), Ragnar Söderbergs Stiftelse (SJE), Knut och Alice Wallenberg Foundation, Sweden (2017–0276, SJE), Swedish Foundation for Strategic Research (FFL18-0049, SJE), the Swedish research council (2017-01874, LOT), Alzheimerfonden (AF930954 LOT, AF940067S SSW), Hjärnfonden (F02018-0325, LOT), Stiftelsen Olle Engkvist byggmästare (194-0675, SSW), KID funding (2016-00150, SSW).

## SUPPORTING INFORMATION

### METHODS AND MATERIALS

#### DNA constructs

A sfGFP150TAG (superfolder GFP) reporter construct with four tandem h7SK-Mma PylT repeats has been described previously (Addgene #140015) [32]. The *M. mazei* PylT/RS expression plasmid B213 was generated by inserting four tandem h7SK-Mma PylT into the unique SpeI site in the published *M. mazei* 4xU6-PylT/RS expression plasmid [18]. All DNA constructs were verified by Sanger sequencing. Sequence of B213 is attached below and available at https://benchling.com/organizations/elsasserlab_group

#### Non canonical amino acid and bioorthogonal labeling

The non canonical amino acid N-ε-[(2-methyl-2-cyclopropene-1-yl)-methoxy]-L-lysine (CpK, SiChem, SC-8017) was used for genetic code expansion. Stock solutions were prepared at 100 mM in 0.2 M NaOH/H2O, 15% DMSO. The GFP150CpK amber suppression reporter was labelled by SPIEDAC with Silicon rhodamine (SiR)-tetrazine (Spirochrome, SC-008). SiR-tetrazine 10 mM stock solution was prepared in DMF.

#### Cell Culture

##### Human induced pluripotent stem cell culture

hiPSCs (CTL07-II iPS) were purchased from the human iPS Core facility at Karolinska Institutet [30]. CTL07-II is a Male cell line, reprogrammed from fibroblasts, with normal 46 XY karyotype. Registration number: 2012/208-31/3.

Coating: 12-well plates were coated with 500 µl laminin-521 (Biolaminina, LN521) diluted 20x in PBS per well at 4°C overnight or at 37°C for 2 hours.

Medium: hiPSCs were cultured in Complete E8 medium (Gibco, A1517001) with Penicillin-Streptomycin (Sigma, P4333-100ml).

Culturing: hiPSCs were washed once with PBS (Sigma, A6964-500ML). 500 µl TrypleSelect (Gibco, 12563029) were added per well in a 12-well plate for 3 minutes at 37°C. Detached cells were transferred into a tube with 1 ml medium and centrifuged for 3 minutes at 300 g. Afterwards cells were resuspended in fresh medium and 10 µM ROCK Inhibitor (Millipore, Y-27632) was added. 75% of the medium was changed every day and cells were passaged every 3-4 days. Cells were imaged in a ZOE Fluorescent Cell Imager (BioRad).

##### Neural stem cell culture

Coating: 12-well plates were first coated with 0.02 mg/ml poly-ornithine (Sigma, #P3655-100MG) in PBS for 30 minutes at 37°C. After washing with PBS twice, culturing plates were incubated with 2-4 µg/ml Laminin 2020 (Sigma, #L2020) in PBS at 4°C overnight or at 37°C for 4 hours.

Medium: To prepare 50 ml of Neural Expansion medium for culturing NSCs, 48.5 ml DMEM/F12+Glutamax (Gibco, #31331-028) were mixed with 500 µl N2 (Gibco, #17504-044), 500 µl Penicillin-Streptomycin, 50 µl B27 (Gibco, #17502-048), 10 ng/ml bFGF (Life Technologies, #CTP0261) and 10 ng/ml EGF (PreproTech, #AF-100-15).

Culturing: The cells were washed once with PBS before passaging. 500 µl Accutase (Sigma, A6964-500ML) were added to each well of the 12-well plates and incubated for 5-8 minutes at 37°C. Detached cells were transferred into a tube with 500 µl PBS and centrifuged for 4 minutes at 300g. The cells were resuspended in PBS and centrifuged on more time at 300g for 4 minutes. The supernatant was aspirated again, and the cells were resuspended in fresh Neural Expansion medium and added into the 12 well plates after removing the Laminin 2020 from the wells. ROCK inhibitor (Millipore, Y27632) was added to a final concentration of 5 µM. The medium was changed every second day and cells were split every 4-5 days.

#### Neurons

Coating: 12-well plates were first coated with 0.02 mg/ml poly-ornithine (Sigma, #P3655-100MG) in PBS for 30 minutes at 37°C. After washing with PBS twice, culturing plates were incubated with 2-4 µg/ml Laminin 2020 (Sigma, #L2020) in PBS at 4°C overnight or at 37°C for 4 hours.

Medium: To prepare 50 ml of medium for culturing neurons, 48.5 ml DMEM/F12+Glutamax (Gibco, #31331-028) were mixed with 500 µl N2 (Gibco, #17504-044), 500 µl Penicillin-Streptomycin and 500 µl B27 (Gibco, #17502-048).

Culturing: Neurons were cultured for 3-4 weeks. The medium was changed every second day and from day 14 on 2-4 µg/ml Laminin 2020 (Sigma, #L2020) was added to the medium.

#### Generation of stable cell lines

Stable cell lines were generated as published [31] with minor adaptations. In short, 100.000 hiPSCs (CTL-07-II iPS) were seeded in a 12-well plate 24 h before transfection. 5 µg DNA in 2:1 ratio (PiggyBac vector : pPBase vector) were diluted in 100 µl OPTI-MEM (Thermofisher, 31985070) before adding LT-1 (Mirus/Kem-EN-Tec-Nordic, MIR 2305). The transfection mix was incubated for 15 minutes to allow formation of transfection complexes, resuspended in 900 µl medium and added onto the cells. In the first step, the synthetase plasmid was integrated, and stable cell lines were generated through selection with 1 µg/ml Puromycin (VWR, CAYM13884-100). After selection the cells were passaged in presence of apoptosis inhibitor CloneR (StemCell, #05888). In the second step the reporter plasmid (Addgene, # 140015) [32], was transfected, and the cells were selected with 200 µg/ml Blasticidin (Invivogen, ant-bl-10p) and 1 µg/ml Puromycin. After selection the cells were again passaged in presence of apoptosis inhibitor CloneR inhibitor (StemCell, #05888).

#### Differentiation of hiPSCs to NSCs and neurons

hiPSCs were differentiated to NSCs with Neural Induction Medium (Gibco, A1647801) according to the manufacturer protocol. In short, 24 h after splitting hiPSCs, at a confluency of around 20%, the medium was completely exchanged to 1 ml of the neural induction medium. Two days later, the medium is exchanged again. After 4 and 6 days, the medium was exchanged again, but 2 ml new medium were added into the 12-well plate. After 7 days of neural induction the cells were split into wells coated with poly-ornithine and L2020. NSCs were differentiated to neurons by removing bFGF and EGF from the medium and increasing B27 (Gibco, #17502-048) to 1:100.

#### Generation of cerebral organoids

Cerebral organoids were generated with help of the STEMdiff™ Cerebral Organoid Kit (StemCell, #08570) according to the manufacturer’s protocol. From day 25 to 40, 0.2 mM CpK was added to the medium.

#### Organoid light-sheet microscopy and immunofluorescence miscroscopy

Living organoids were imaged on day 40 with the Zeiss Light Sheet Z.1. Organoids were mounted in 1% low-melt agarose (LMA) in a glass capillary and the entire 3D volume of the organoid was imaged with a pixel width and height of 0.644 µm and a voxel depth of 2 µm. Afterwards organoids were fixed according to the previously published protocol from Lancaster and Knoblich [9]. In short, organoids were washed with PBS and fixed in 4% Formaldehyde (Thermofisher, 28906) at 4°C for 15 minutes. Afterwards organoids were washed 3 times with PBS and embedded in 30% sucrose (w/v) (Sigma, S0389-500G) solution at 4°C overnight. The next day, organoids were placed into a 7.5% gelatin/10% sucrose (w/v) embedding solution at 4°C, until the solution polymerizes. Organoids were frozen in a bath of Isopentane and stored at −80°C. Before Cryosectioning, the organoids were warmed to −20°C overnight. 20 µm slices of the organoids were cut via cryosectioning and washed with PBS-T at 37°C for 10 minutes to fully remove gelatin from slides. Afterwards slides were blocked in 5% Normal Donkey Serum in PBS-T for 1h and stained with different antibodies overnight at 4°C. Description of the antibodies and their dilution in 5% BSA in PBS-T can be found in the table below. The next day, slides were washed 3 times with PBS-T before the secondary antibody was added. After 2 hours incubation at room temperature, slides were washed 3 times in PBS-T for 30 minutes each. After cells were dried for 5 minutes Duolink® In Situ Mounting Medium with DAPI (Sigma, DUO82040) was added. Slides were imaged using a Zeiss AiryScan800 laser scanning confocal microscope. Lasers and filters were chosen to illuminate at fluorophore excitation maximum. Images were processed using the software ImageJ. Illumination and gain settings were kept constant for all experiments.

| Antibody | Dilution | Supplier |
| --- | --- | --- |
| CTIP2 | 1:2000 | Abcam (ab18456) |
| SOX2 | 1:2000 | Rndsystems (MAB2018) |
| GFP | 1:1000 | Chromotek (PABG1-100) |
| MAP2 | 1:1000 | Millipore (AB5622) |
| Donkey anti-Rabbit IgG, Secondary Antibody, Alexa Fluor 488 | 1:500 | Life Technologies (A-21200) |
| Donkey anti-Mouse IgG, Secondary Antibody, Alexa Fluor Plus 647 | 1:500 | Life Technologies (A-21236) |

#### Immunofluorescence

Cells were washed with PBS and fixed in 4% Formaldehyde (Thermofisher, 28906) at 4°C for 15 minutes. Afterwards cells were washed 3 times with PBS and permeabilized with 0.1% Triton X-100 for 15 minutes. Cells were blocked in 0.1 % BSA in 0.05% Triton-X-100 in PBS solution for 1 hour, before incubation with primary antibodies at 4°C overnight. The source of the antibodies and their dilution in 0.1 % BSA in 0.05% Triton-X-100 in PBS can be found in the table below. The next day, cells were washed 3 times with PBS. Secondary antibodies were added for 2 hours, before the cells were washed again 3 times with PBS. After cells were dried for 5 minutes Duolink® In Situ Mounting Medium with DAPI (Sigma, DUO82040) was added. Slides were imaged using a Zeiss AiryScan800 laser scanning confocal microscope. Lasers and filters were chosen to illuminate at fluorophore excitation maximum. Images were processed using the software ImageJ. Illumination and gain settings were kept constant for all experiments. For the cell analysis, the software CellProfiler was used. To quantify the differentiation markers (SOX2, OCT4, NESTIN and MAP2) mean fluorescence intensity in each image, corrected for the number of cells, were measured. The mean intensity of GFP expression was measured for each cell.

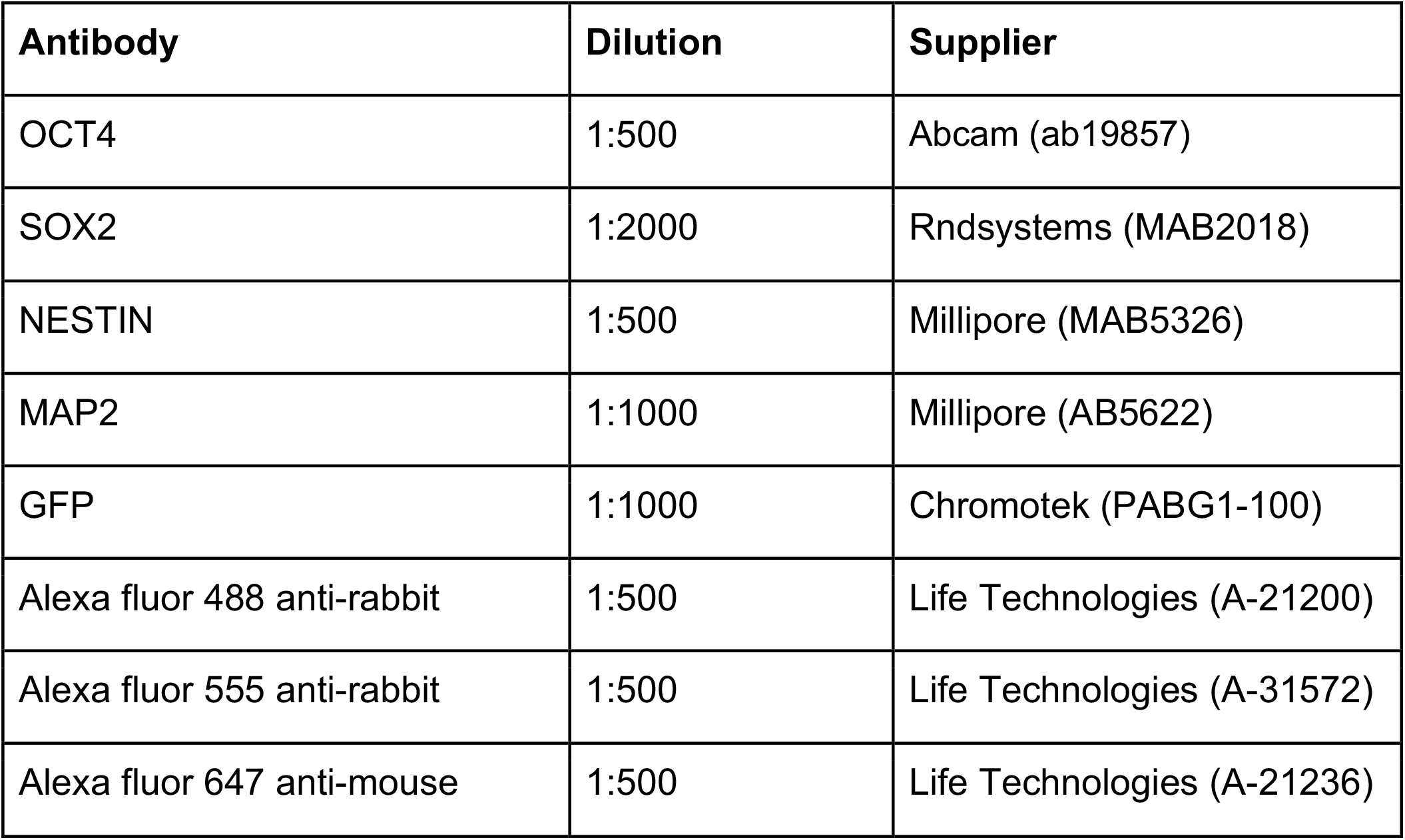

#### Bioorthogonal SPIEDAC labeling of neurons

Neurons were washed 3 times with medium and incubated for 2 h to remove excessive CpK. Afterwards the cells were cultured in the presence of 0.5 µM SiR-tetrazine (Spirochrome, SC008) for 15 min, before the cells were again 3 times washed and incubated for 1 h. After a last washing step, it was continued as described in the method section Immunofluorescence.

#### Western blots

Cells were lysed in N-Ripa Buffer (10 mM Tris-HCl (pH 7.6), 1 mM EDTA, 0.1% sodium dodecyl sulfate, 0.1% sodium deoxycholate, 1% Triton X-100, 5% Glycerol, 140 mM NaCl, 1x protease inhibitor) and sonicated for 20 cycles (30 sec on/30 sec off). Protein concentration was determined by BCA assay (Thermofisher, 23227). For lysate labeling 1 µM SiR-tetrazine (Spirochrome, SC008) was added to the lysate for 5 min. SDS PAGE sample buffer was added and samples were boiled briefly at 95°C. 25 μg protein were loaded in each well. Samples were loaded on 4–20% polyacrylamide Bis-Tris gels (BioRad, #4561096) and exposed for in-gel fluorescence at 460nm and 630 nm. The protein was transferred to nitrocellulose membranes and probed with FLAG-Antibody and GAPDH antibodies and imaged with the GE Healthcare ImageQuant LAS 500.

| **Antibody** | **Dilution** | **Supplier** |
| --- | --- | --- |
| FLAG | 1:5000 | Sigma (A8592) |
| GAPDH | 1:10000 | Millipore (AB2302) |
| GFP | 1:500 | ChromoTek (pABG1-100) |
| OCT4 | 1:500 | Abcam (ab19857) |
| H3 | 1:15000 | Active Motif (39763) |
| anti-mouse HRP | 1:10000 | Biorad (Cat#1721011) |
| anti-rabbit HRP | 1:10000 | Biorad (Cat#1721019) |
| anti-chicken IgY (H+L) HRP | 1:10000 | Invitrogen (A16054) |

## SUPPLEMENTARY FIGURES

**Supplementary Figure S1:**
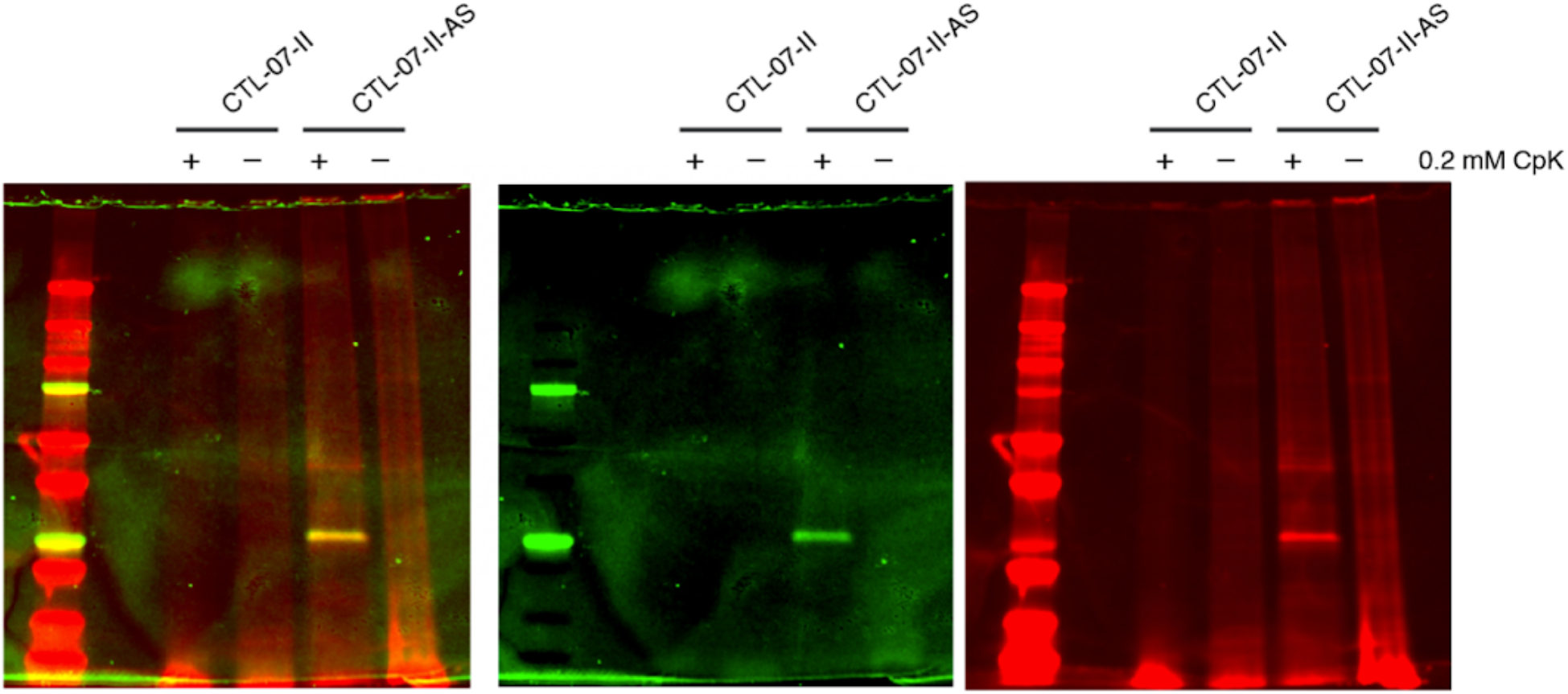
Bioorthogonal SPIEDAC labeling of lysate from CTL07-II hiPSCs and CTL07-II-AS hiPSCs. Cells were cultured in the presence of 0.2 mM CpK for 24 h (+) or without CpK (-). Cell lysate was incubated with SiR-Tetrazine for 5 min, before it was separated by SDS-PAGE and imaged for in-gel fluorescence.

**Supplementary Figure S2:**
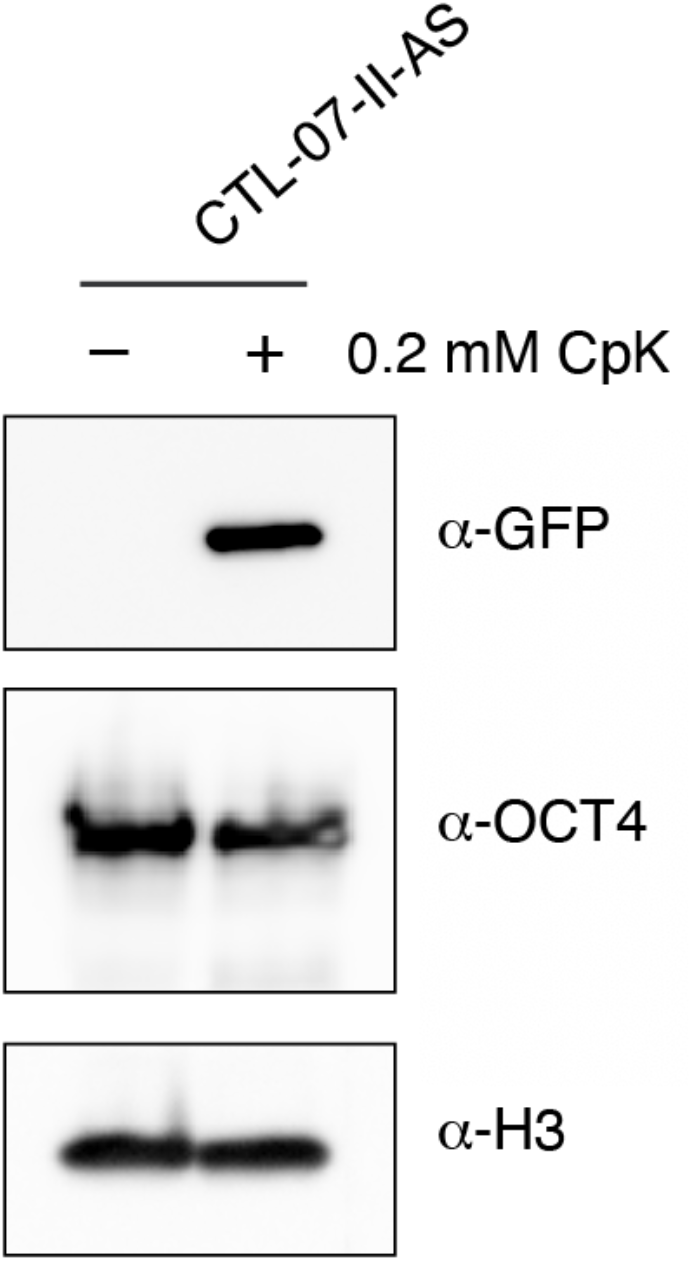
Western blots from cell lysates of CTL07-II-AS showing expression of GFP when cultured in the presence of 0.2 mM CpK for 24 h. OCT4 was expressed in cells cultured in the presence and absence of CpK. H3 was used as loading control.

**Supplementary Figure S3:**
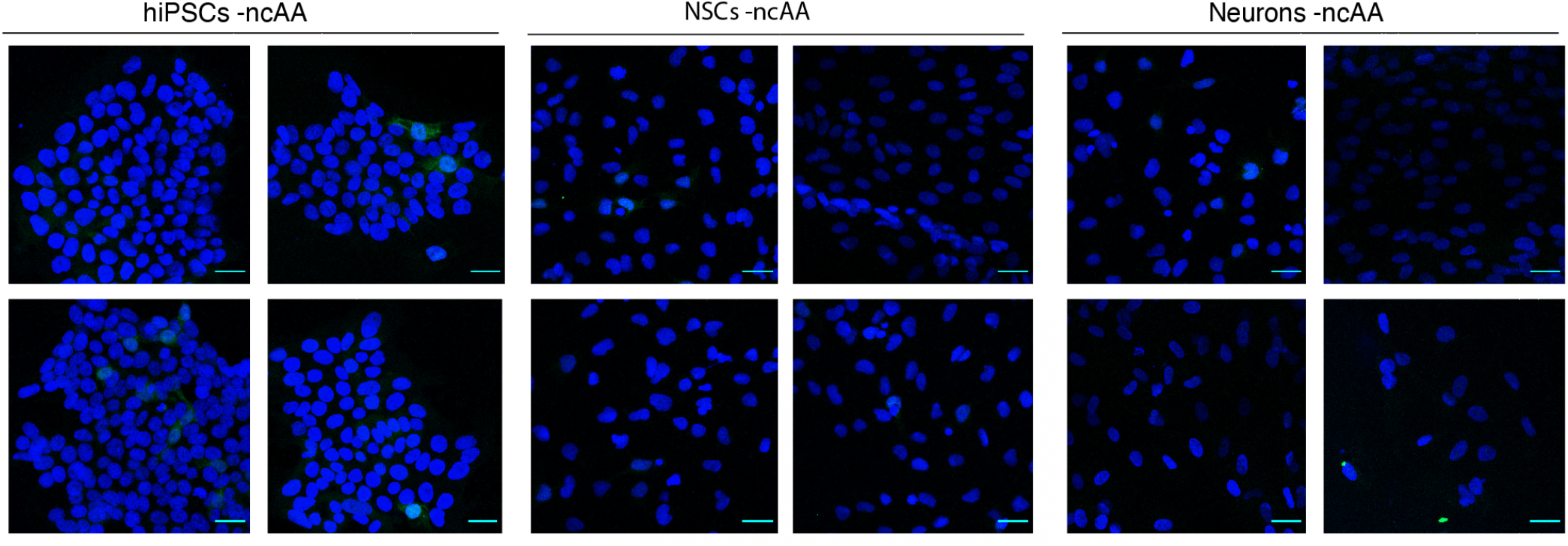
Four representative images from immunofluorescence live cell microscopy of CTL07-II-AS hiPSCs, derived NSCs and neurons showing expression of GFP (green) and DAPI (blue), cultured in the absence of ncAA.

**Supplementary Figure S4:**
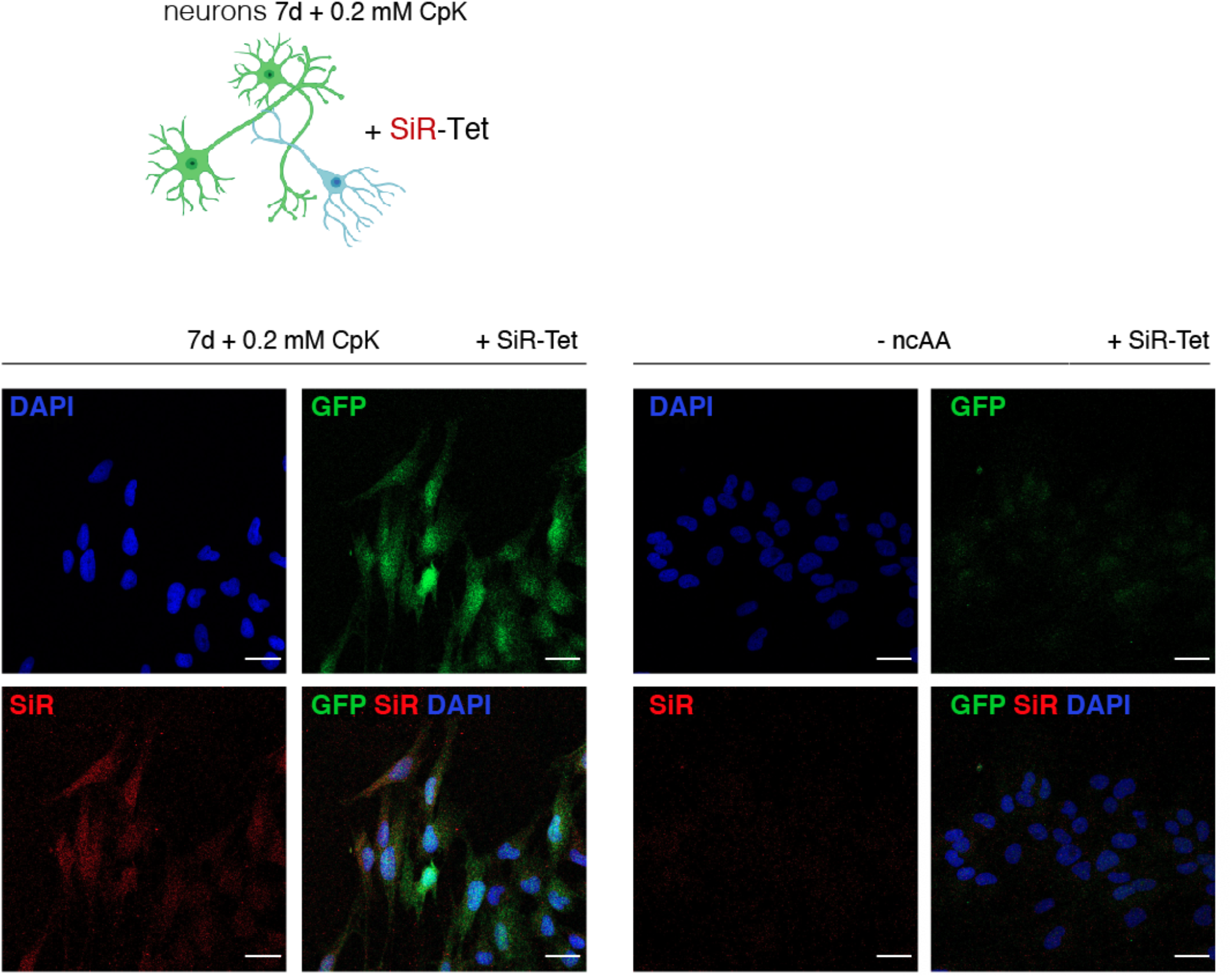
Representative images of SPIEDAC labeled 3 week old neurons. Neurons were cultured in the presence or absence of 0.2 mM CpK for 7 days, before cells were incubated in 0.5 µM SiR-Tetrazine dye for 15 min before fixation. GFP expression is shown in green, SiR-Tetrazine labeling in red and DAPI in blue. Scale bars correspond to 30 µm.

**Supplementary Figure S5:**
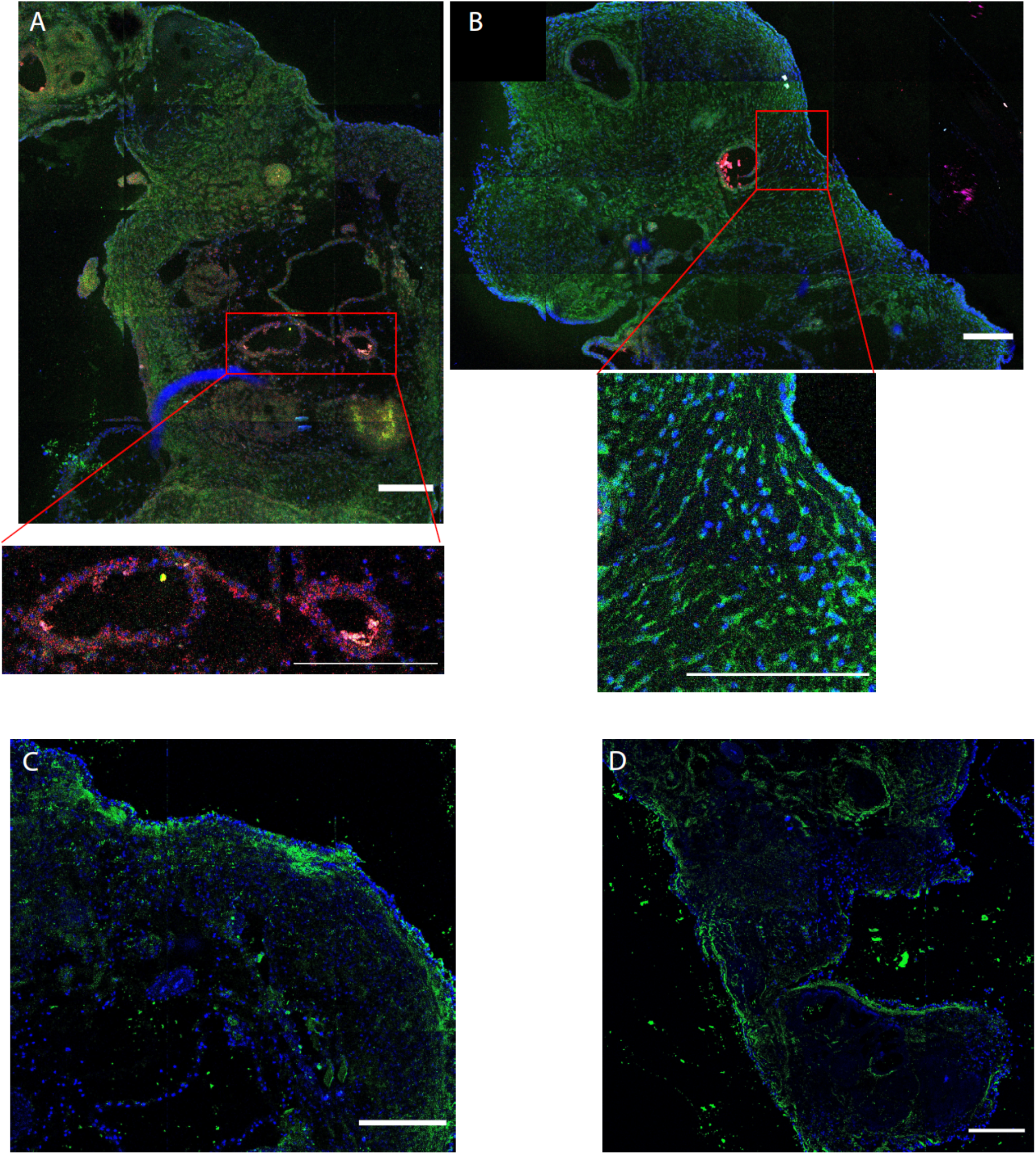
Characterization of cerebral organoids generated from CTL-07-II-AS hiPSCs and differentiated for 40 days. A) DAPI (blue) and SOX2 (red) co-staining highlights rosette-like progenitor compartments around the ventricles. B) DAPI (blue) and MAP2 (green) show successful differentiation to neurons. C-D) DAPI (blue) and CTIP2 (green) staining shows deep-layer cortical neurons on the surface of the cerebral organoids.

**Supplementary Figure S6.**
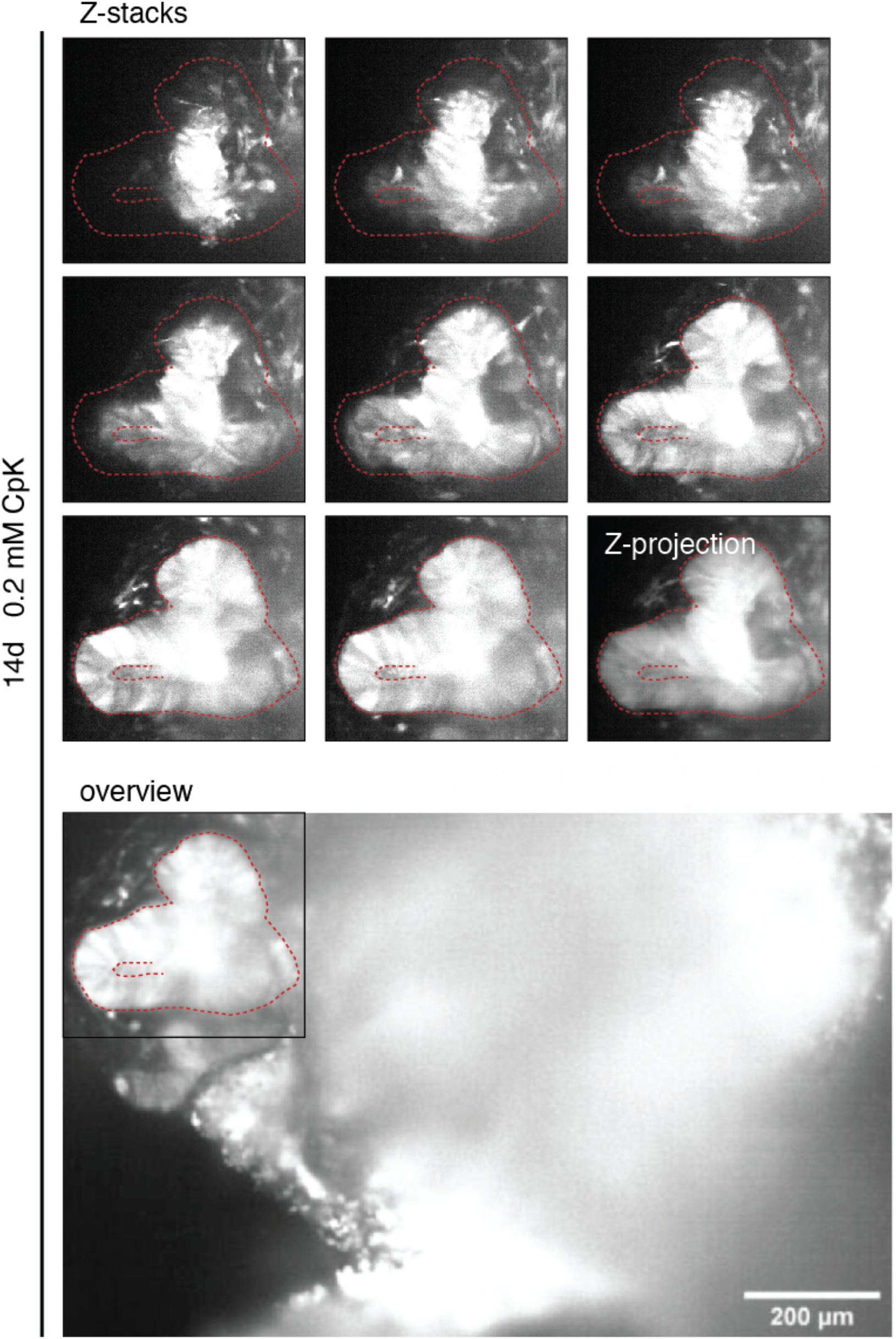
Stable amber suppression in cerebral organoids. Live cell light-sheet microscopy of cerebral organoids to detect sfGFP expression in organoids cultured with 0.2 mM CpK Scale bars correspond to 200 µm. Luminal neuroepithelial structure with high GFP fluorescence is highlighted in red. See also entire Z-stack (Movie 4) and 3D reconstruction (Movie 5).

**Supplementary Movie 1: Z-stack, GFP fluorescence, cerebral organoid**

Live cell light-sheet microscopy (GFP channel) of a cerebral organoid cultured with 0.2 mM CpK.

**Supplementary Movie 2: Z-stack, GFP fluorescence, tip of cerebral organoid**

Live cell light-sheet microscopy (GFP channel) of the tip of a cerebral organoid cultured with 0.2 mM CpK.

**Supplementary Movie 3: 3D projection, GFP fluorescence, tip of cerebral organoid**

3D reconstruction from live cell light-sheet microscopy (GFP channel) of the tip of a cerebral organoid cultured with 0.2 mM CpK.

**Supplementary Movie 4: Z-stack, GFP fluorescence, luminal cluster**

Live cell light-sheet microscopy (GFP channel) of a luminal rosette within a cerebral organoid cultured with 0.2 mM CpK.

**Supplementary Movie 5: 3D projection, GFP fluorescence, luminal cluster**

3D reconstruction from live cell light-sheet microscopy (GFP channel) of a luminal rosette within a cerebral organoid cultured with 0.2 mM CpK.

## UNPROCESSED WESTERN BLOT IMAGES

Western blots from cell lysates of CTL07-II-AS showing expression of FLAG in comparison to the parental CTL07-II cells.

**Figure 2.**
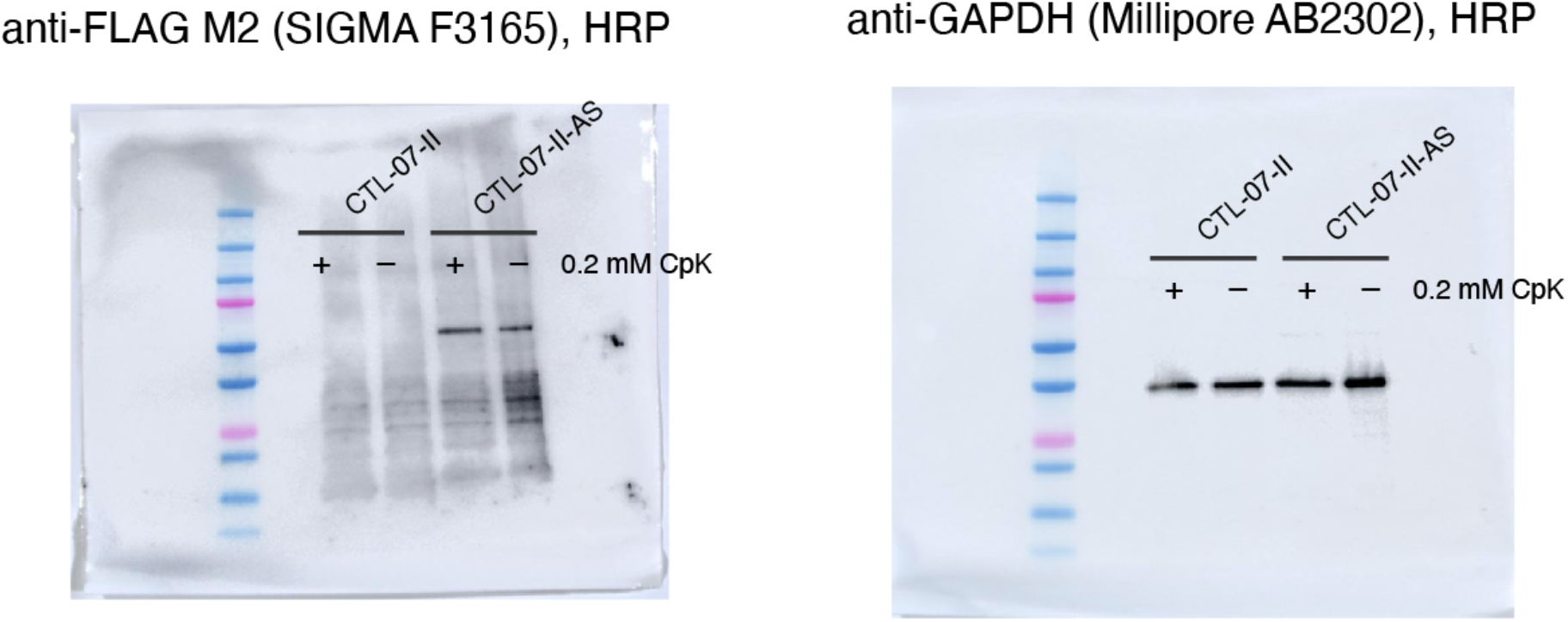

**Supplementary Figure 2.**
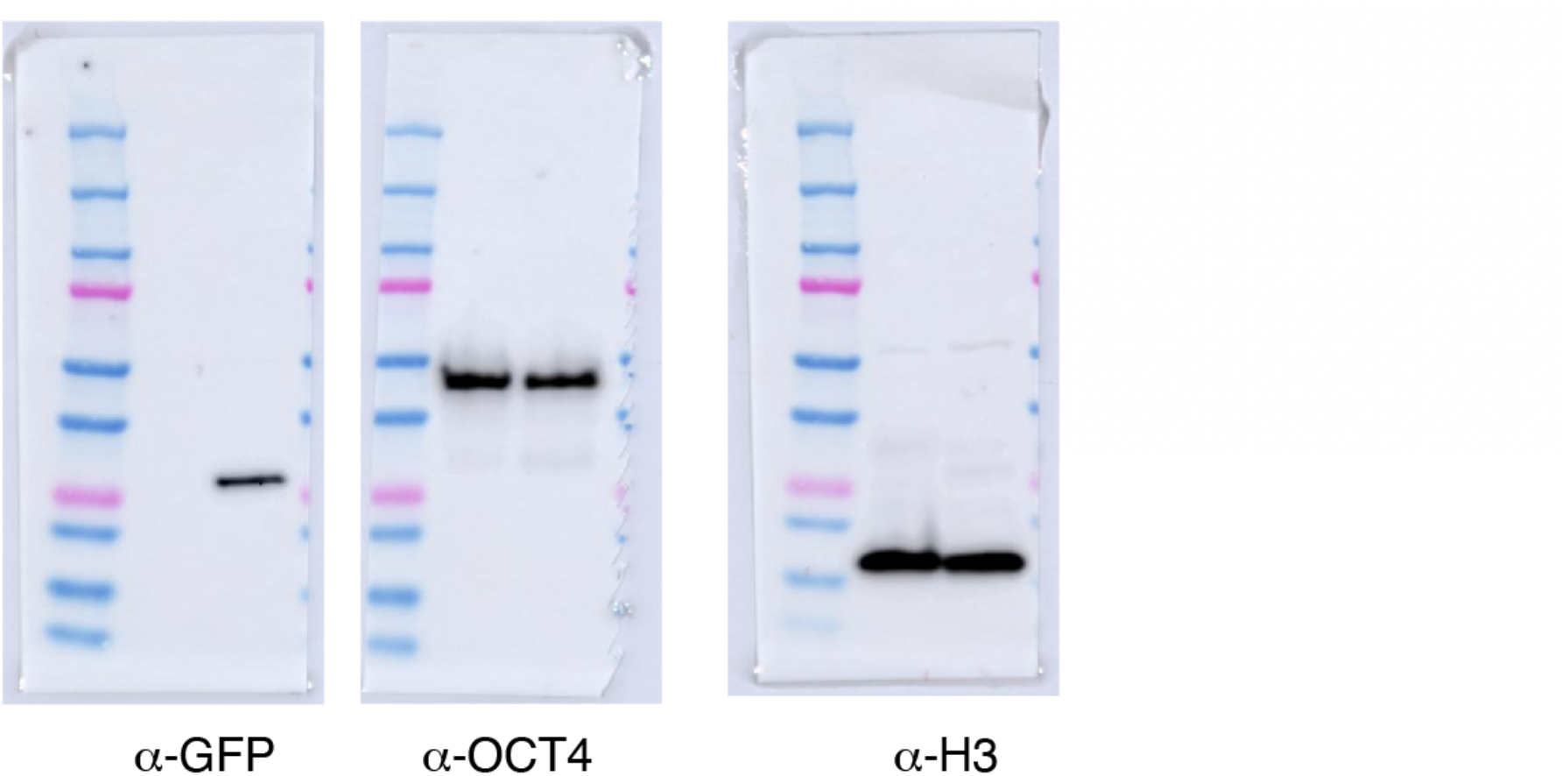

Western blots from cell lysates of CTL07-II-AS showing expression of GFP when cultured in the presence of 0.2 mM CpK for 24 h. OCT4 was expressed in cells cultured in the presence and absence of CpK. H3 was used as loading control.

## Plasmid Sequence B213

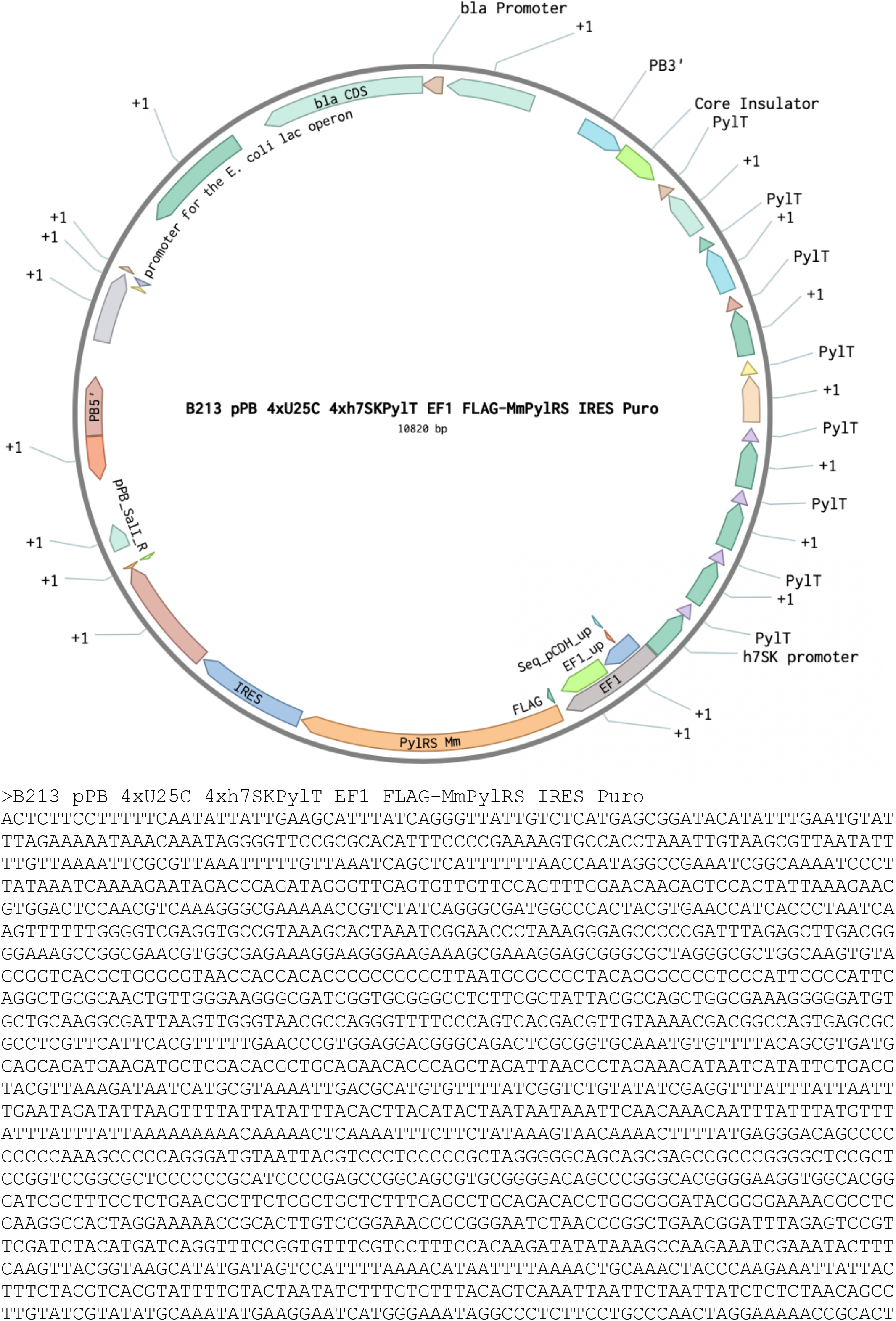
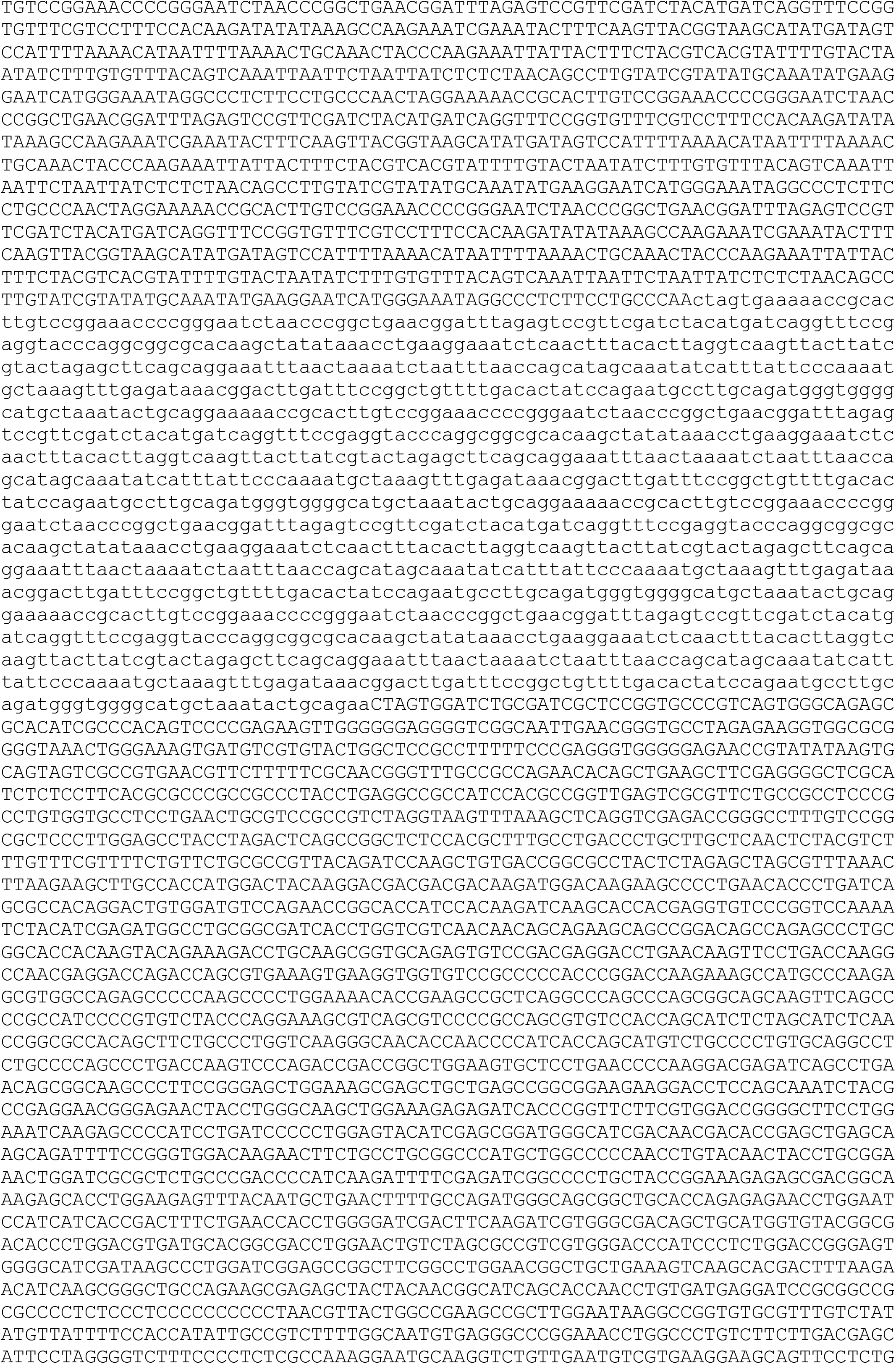
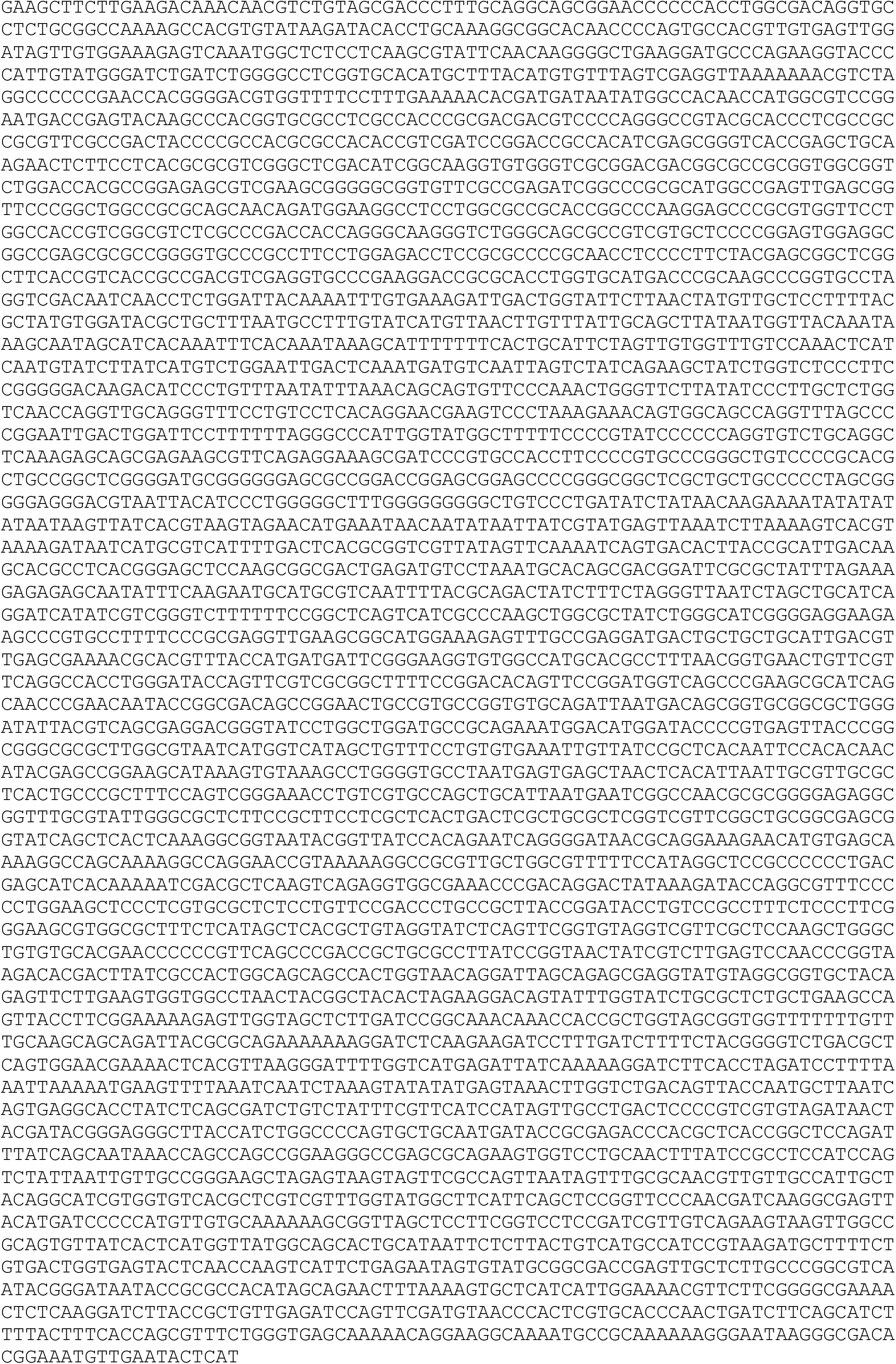

